# The autism-associated *Meis2* gene is necessary for cardiac baroreflex regulation in mice

**DOI:** 10.1101/2022.09.14.507907

**Authors:** J Roussel, R Larcher, P Sicard, P Bideaux, S Richard, F Marmigère, J Thireau

**Author notes:** Corresponding authors. Corresponding authors: Marmigère Frédéric, Institut de Génomique Fonctionnelle de Lyon (IGFL), École Normale Supérieure de Lyon, CNRS, Lyon, France. Mail, Thireau Jérôme, Phymedexp, Université de Montpellier, Inserm, CNRS, CHRU de Montpellier, Montpellier, France. Mail. Equal contributions (F Marmigère, J Thireau).

## Abstract

Recent understanding of Autism Spectrum Disorder (ASD) showed that peripheral primary mechanosensitive neurons involved in touch sensation and central neurons affected in ASD share transcriptional regulators. Mutant mice for ASD-associated transcription factors exhibit impaired primary tactile perception, and restoring those genes specifically in primary sensory neurons rescue some of the anxiety-like behavior and social interaction defects.

Interestingly, peripheral mechanosensitive sensory neurons also project to internal organs including the cardio-vascular system, and an imbalance of the cardio-vascular sympatho-vagal regulation is evidenced in ASD and intellectual disability. ASD patients have decreased vagal tone, suggesting dysfunction of sensory neurons involved in cardio-vascular sensing.

In light of our previous finding that the ASD-associated *Meis2* gene is necessary for normal touch neurons development and function, we investigated here if its inactivation in mouse peripheral sensory neurons also affects cardio-vascular sympatho-vagal regulation and baroreflex. Combining echocardiography, pharmacological challenge, blood pressure monitoring and heart rate variability analysis, we found that *Meis2* mutant mice exhibited a blunted vagal response independently of any apparent cardiac malformation. These results suggest that defects in primary sensory neurons with mechanosensitive identity could participate in the imbalanced cardio-vascular sympatho-vagal tone found in ASD patients, reinforcing current hypotheses on the role of primary sensory neurons in the etiology of ASD.

## Introduction

Autism Spectrum Disorder (ASD) is the consequence of a neurodevelopmental defect affecting different nervous system structures and characterized by many diverse phenotypic manifestations including aberrant social interactions, repetitive behaviors and restrictive interest. In addition, 90% of ASD patients are estimated to present sensory processing deficits, an inability to elaborate appropriate behavioral responses due to impaired sound, touch and sight perception^1^. This defective sensory perception can lead to an altered functional “vagal brake” associated with a defective behavioral flexibility to stress^2^.

A large number of genes have been associated with ASD and are believed to be involved in various stages of building neuronal architecture, from neurogenesis to neurites outgrowth, synaptogenesis and synaptic plasticity^3–6^. The diverse cellular expression and functions of ASD-associated genes across brain regions and neuronal cell types is reflected in the wide range of common and divergent phenotypic outcomes. Consequent to this genetic diversity, phenotypic characterization of the syndrome has often proven difficult, resulting in inconsistent conclusions. For instance, despite a paucity of information and conflicting findings in the literature, imbalance between the sympathetic and parasympathetic branches of the autonomic nervous system is commonly observed in ASD patients^7–19^. Overall, these studies point to a lower autonomic nervous system activity suggested to likely result from a decreased parasympathetic activity. More strikingly, characterization of autonomic activity in Rett syndrome, one of the most characterized ASD-related disorder, illustrates the diversity of the phenotypic manifestation of the vagal imbalance. Whereas some studies report a vagal imbalance with increased LF/HF ratio and HF component, others report a decreased cardiac baroreceptor sensitivity and cardiac vagal tone^7,20–24^. In the first case, it was suggested that individuals suffering Rett syndrome have an increased sympathetic activity that is not counterbalanced by vagal tone, whereas in the second case, the authors concluded that Rett patients exhibit a low cardiovascular parasympathetic tone but a normal sympathetic activity. Nevertheless, in line with current emerging hypothesis of the role of primary sensory neurons in the etiology of ASD, these observations raise the possibility that peripheral neurons in general and peripheral sensory neurons in particular are defective in some ASDs.

Recent advances in the understanding of ASD suggest that centrally affected neurons in ASD and peripheral touch mechanosensitive sensory neurons of the Dorsal Root Ganglia (DRG) share specific transcriptional programs regulating late neuronal differentiation^25,26^. These touch neurons express several of the ASD-linked genes, and mutant mouse models for ASD exhibit primary sensory deficits. Specific inactivation of ASD-associated genes in the peripheral somatosensory system recapitulated some ASD symptoms such as altered cognitive and social behavior. Conversely, tissue-specific re-introduction of those genes in full knockout models not only rescued the normal functioning of primary touch neurons, but also some of the anxiety-like and altered social behaviors^25,26^. Thus, specific inactivation of ASD-associated genes allows the uncoupling of complex and intermingled ASD-associated symptoms.

Among the genes recently associated with ASD, the transcription factor (TF) *MEIS2* is a strong candidate to participate to autonomic regulation of cardiac rhythm^27–29^. *MEIS2* is a member of the *MEIS* (Myeloid Ecotropic viral Insertion Site) family of homeobox TFs that belongs to the Three Amino-acid Loop Extension (TALE) family. These TFs are involved in the embryonic development of a plethora of organs and cell types, in particular in the nervous system^30,31^. In mice, Meis TFs are also strongly linked to heart embryonic development and postnatal functions^32–36^, and in humans *MEIS2* haploinsufficiency causes severe neurodevelopmental defects with intellectual disability and ASD-like behavioral abnormalities, cleft palate and heart defects^27–29^. Combining mouse genetic and Heart Rate Variability (HRV) approaches, we previously showed that, independently of any heart morphological defects, specific *Meis1* inactivation in mouse developing sympathetic resulted in a severe cardiac chronotropic incompetence eventually leading to sudden cardiac death^31^. This phenotype was attributed to a failure by sympathetic neurons to complete distal innervation of target organs, including the heart. Thus, combining conditional gene ablation in mice, HRV analysis and pharmacological testing of heart rate adaptation to blood pressure changes offers a powerful workflow to disentangle the mechanisms leading to cardiac dysautonia.

More recently, we found that *Meis2* inactivation in postmitotic peripheral sensory neurons dictates comparable phenotypes for DRG touch neurons with incomplete distal innervation, impaired electrophysiological responses to mechanical stimuli and reduced touch sensation^36^. In the present study, because *Meis2* is essential for normal heart development, we took advantage of the same mouse strain as used previously to investigate, independently of cardiac malformations, if *Meis2* targeted inactivation in peripheral sensory neurons also recapitulates some of the cardiac sympatho-vagal imbalance phenotypes reported in ASD studies. Our results showed that indeed *Meis2* expression in somatosensory neuron*s* is indispensable for functional adaptation of cardiovascular parameters. These mutant mice exhibited increased sinus rhythm variability and modified sympatho-vagal index together with altered cardio-inhibitory reflex (cardiac baroreflex) independently of any cardiac morphological and contractile defects. These results are consistent with the decreased cardiac baroreceptor sensitivity reported in ASD, and the decreased cardiac vagal tone and cardiac sensitivity to baroreflex in Rett patients, suggesting that suppressing *Meis2* function in late differentiating peripheral neurons recapitulates some of ASD symptoms.

## Results

### Meis2 mutant mice do not present any morphological or contractile heart defects

The mouse strain used here was Isl1^+/CRE^::Meis2^LoxP/LoxP^ in which the 8^th^ homeobox-containing exon was flanked by LoxP sites^36^. Both in mouse and human, *Meis2* mutations causes severe developmental anomalies of the heart. Humans carrying heterozygous *MEIS2* missense mutations or 15q14 microdeletion involving *MEIS2* present a triad of cleft palate, atrial or ventricular septal heart defects, and developmental delay^27–29^. In another *Meis2*-null mouse strain, an incomplete septation of the outflow tract known as persistent truncus arteriosus was reported, and specific Meis2 ablation in cardiac neural crest precursors led to a defective heart outflow tract^33^. Islet1 (Isl1) is also expressed in distinct cardiovascular lineages^37^ raising the possibility that CRE recombination in Isl1^+/CRE^::Meis2^LoxP/LoxP^ mice results in heart malformations. To ascertain whether the Isl1^+/CRE^::Meis2^LoxP/LoxP^ strain allows investigating cardiac autonomic function independently of heart malformations, we morphologically and functionally characterized the adult heart in WT, Isl1^+/CRE^ and Isl1^+/CRE^::Meis2^LoxP/LoxP^ mice using Doppler echocardiography (Figure 1). Isl1^+/CRE^ mice were included as an additional control group to avoid misinterpretation due to Isl1 heterozygosity. Investigations of the parasternal long and short axis views (Figure 1) did not reveal any cardiac malformation. Thicknesses of the septum (IVS;d, IVS;s, Figure 1A) and of the left ventricular posterior wall in diastole and in systole periods (LVPW;d and LVPW;s; Figure 1B) were identical in all groups. The left ventricle internal diameters were also indistinguishable among groups whatever the cardiac period (LVID;d and LVID;s; Figure 1C). The ejection fraction (EF) and fractional shortening (FS), which are used as the conventional contractile function indexes, were also similar in WT, Isl1^+/CRE^ and Isl1^+/CRE^::Meis2^LoxP/LoxP^ mice (Figure 1D). Finally, heart diastolic performances assessed by measuring left ventricle filling waves in standard 4 cavities view (E/A ratio, Figure 1E) and Aortic flow Velocity Time Integral (Ao VTI, Figure 1F) did not show any difference attesting for normal hemodynamic parameters and contractile performances, and suggesting that the outflow tract was not affected in Isl1^+/CRE^ and Isl1^+/CRE^::Meis2^LoxP/LoxP^ mice (Figure 1G). To conclude, these observations clearly exclude cardiac malformations or remodeling that may alter cardiac function, and allow thus investigating cardiac autonomic regulation independently.

**Figure 1:**
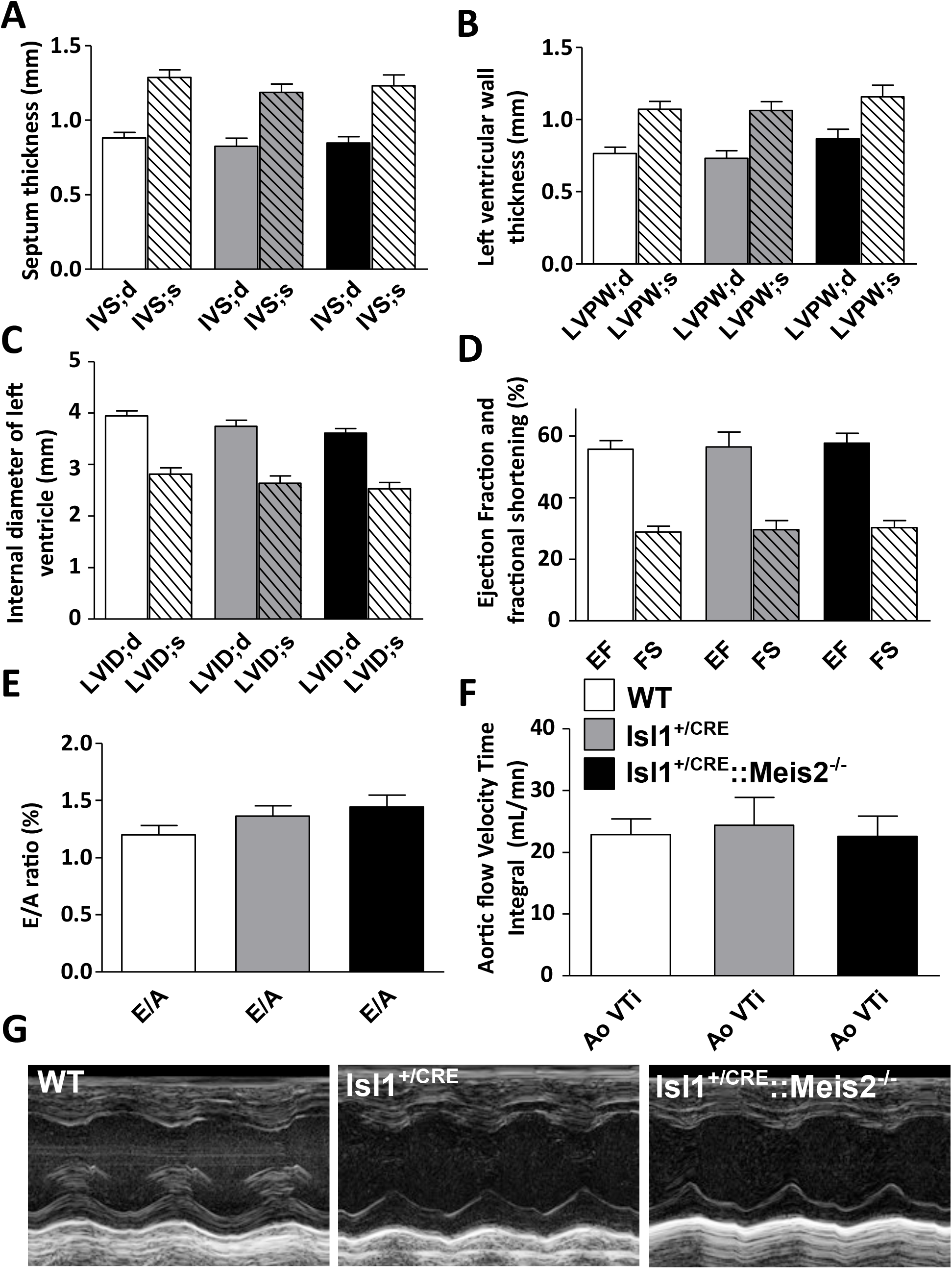
Isl1^+/CRE^::Meis2^LoxP/LoxP^ mice are devoid of morphological and contractile heart defects. Morphologic and left ventricular function parameters assessed by Doppler echocardiography in WT, Isl1^+/CRE^ and Isl1^+/CRE^::Meis2^LoxP/LoxP^ mice. Measures of IVS (**A**), LVPW (**B**), LVID (**C**), EF and FS (**D**) showed no difference between WT, Isl1^+/CRE^ and Isl1^+/CRE^::Meis2^LoxP/LoxP^ mice. Peak early (**E**) and late atrial contraction (**A**) mitral inflow waves velocities were measured and the E/A ratio was calculated (**E**) as the ascending aortic blood flow (**F**). Abbreviations: IVS= thickness of the interventricular septum during diastole (d) and systole (s); LVPW= thickness of the posterior wall of the left ventricle during diastole (d) and systole (s); LVID= left ventricular internal diameter during diastole (d) and systole (s); EF (%)=Ejection Fraction in M-mode; FS (%)=Fractional Shortening in M-mode. Pulsed-wave Doppler of the ascending aortic blood flow was recorded permitting measurements of the velocity time integral (AoVTI). n=9 to 11 mice in each group.

### Meis2 mutant mice exhibit increased sinus rhythm variability and modified sympatho-vagal index

We next characterized cardiac electrophysiological activity using telemetric electrocardiogram recording in *Meis2* mutant (Isl1^+/CRE^::Meis2^LoxP/LoxP^) and control (WT and Isl1^+/CRE^) mice. Using the telemetric system allows long-term recording of ECG on non-sedated and unrestrained mice. ECG analyses showed that the 3 strains exhibited comparable electrophysiological characteristics in baseline conditions (Figure 2A and B; n=11, 12, 11 in WT, Isl1^+/CRE^ and Isl1^+/CRE^::Meis2^LoxP/LoxP^, respectively). The mean values of the ventricular cycle length (RR in ms), the PR interval (interval between the onset of atrial depolarization until the beginning of the onset of ventricular depolarization in ms), the QRS (depolarization time of the right and left ventricles in ms) and of QT duration (time to depolarization-repolarization of ventricles in ms) (Figure 2B) were identical in all groups suggesting comparable cardiac conduction and depolarization/repolarization activities. No ectopic atrial or ventricular arrhythmia were detected in none of the 3 genotypes. By contrast, we observed a large sinus rhythm variability when *Meis2* was inactivated (Figure 2C and D). Indeed, as showed by the SDNN assessing the total beat to beat variability of normal sinus beat, Isl1^+/CRE^::Meis2^LoxP/LoxP^ presented an increased variability (*p=0*.*001* and *p=0*.*002 vs* WT and Isl1^+/CRE^ mice respectively), completed by a non-linearity to RR interval (R^2^=0.17) when compared to WT (R^2^=0.88) and Isl1^+/CRE^ (R^2^=0.87) mice (Figure 2D). This could reflect a dysregulation of spontaneous beat-to-beat variability induced by autonomic pathways. We further assessed HRV by spectral analysis using Fast Fourier Transform (FFT) (Figure 3). Low frequency (LF) was non-significantly decreased in Isl1^+/CRE^::Meis2^LoxP/LoxP^ mice using Anova test (*p=0*.*3679*, n=9 in each group, Figure 3A). High frequencies (HF) were similar in all groups (*p=0*.*401*, n=9 in each group; Figure 3B). However, the LF/HF ratio was significantly decreased in Isl1^+/CRE^::Meis2^LoxP/LoxP^ mice compared to WT mice (*p=0*.*016*, n=9), and no difference was observed between WT and Isl1^+/CRE^ mice (Figure 3C). These results obtained by spectral analysis suggest an alteration in the spontaneous control of cardiac rhythm by the autonomic system. Altogether, the large variability of sinus rhythm, the increase in SDNN and its non-linearity to RR, and the decrease of LF/HF ratio suggest an asymptomatic modification of spontaneous beat-to-beat adaptation in *Meis2* mutant mice and an overall decrease in sympatho-vagal activity.

**Figure 2:**
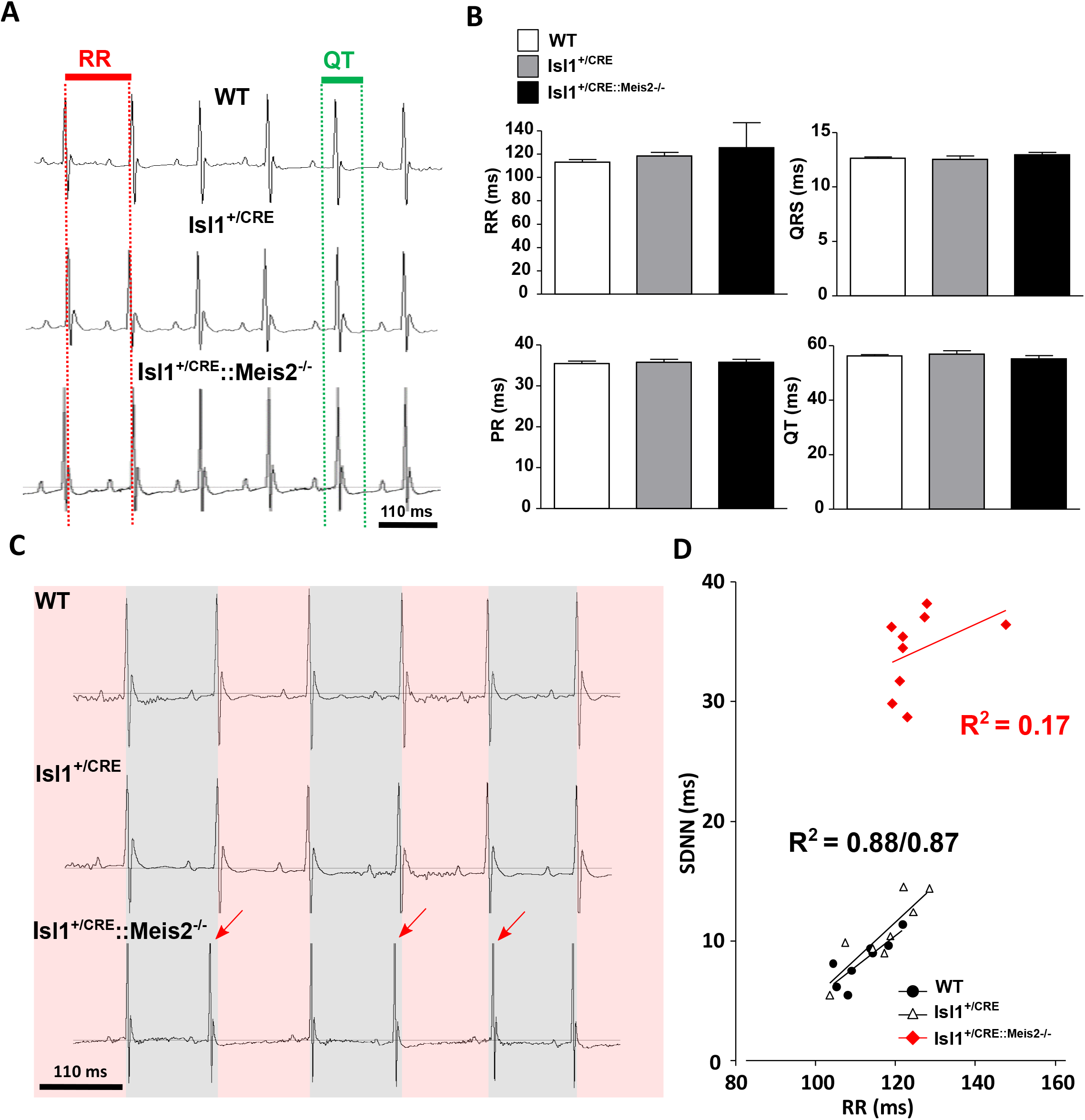
Increased sinus rhythm variability in Isl1^+/CRE^::Meis2^LoxP/LoxP^ mice. Telemetric recording of ECG in vigil WT, Isl1^+/CRE^ and Isl1^+/CRE^::Meis2^LoxP/LoxP^ mice in basal conditions. **A**) Typical ECG traces for WT, Isl1^+/CRE^ and Isl1^+/CRE^::Meis2^LoxP/LoxP^ **B**) Graphs showing the RR, PR, QRS and QT durations in the three groups of mice. No difference was observed between genotypes. n=11 to 12 mice in each group. **C**) Typical ECG traces of sinus variability monitored in WT, Isl1^+/CRE^ and Isl1^+/CRE^::Meis2^LoxP/LoxP^ mice. **D**) Graph showing the correlation between SNDDN and RR in the 3 groups of mice. n=7 to 9 mice in each group. SDNN=Standard Deviation of normal to normal beat intervals.

**Figure 3:**
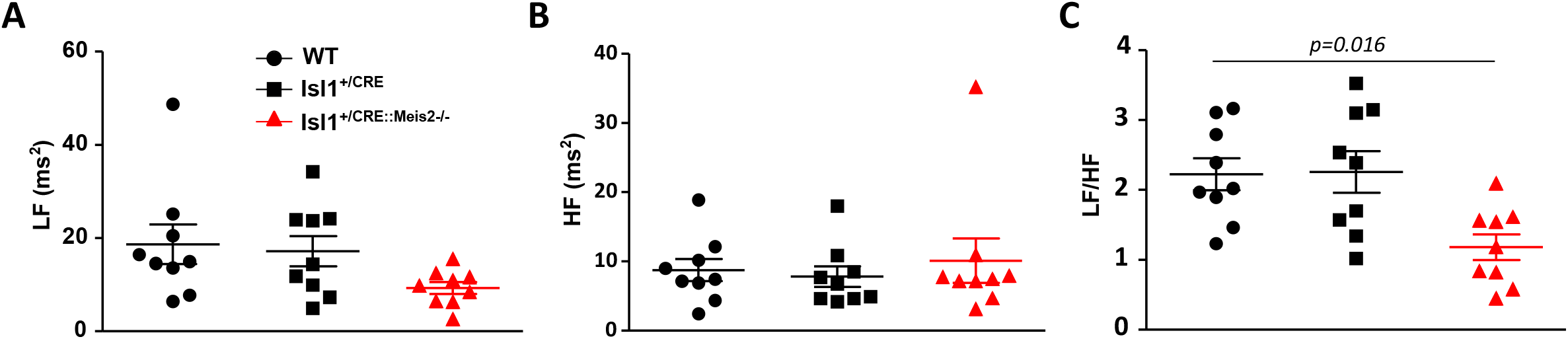
Decreased sympatho-vagal activity in Isl1^+/CRE^::Meis2^LoxP/LoxP^ mice. HRV analysis of WT, Isl1^+/CRE^ and Isl1^+/CRE^::Meis2^LoxP/LoxP^ vigil mice. Low frequency band (LF) (A), High frequency band (B) and sympatho-vagal index LF/H (C) were obtained at baseline by fast Fourier transform and revealed a decrease of sympatho-vagal activity on heart rhythm. n=7 to 9 mice in each group. \**p≤0*.*05* to WT.

### Meis2 mutant mice present non-typical heart rate adaptation with a blunt cardio-inhibitory reflex

To more clearly unmask the autonomic imbalance in Isl1^+/CRE^::Meis2^LoxP/LoxP^ mice, we challenged conscious transmitter-implanted mice with reference drugs. These drugs are well-known to induce fast hemodynamic changes that in turn activate cardiac rhythm adaptation reflexes^38–40^. After nitroprusside injection WT, Isl1^+/CRE^ and Isl1^+/CRE^::Meis2^LoxP/LoxP^ mice presented a reflex increase in heart rate (*p<0*.*001*, n=6, n=6, n=7 for WT, Isl1^+/CRE^ and Isl1^+/CRE^::Meis2^LoxP/LoxP^ animals respectively; Figure 4A). By contrast, Isl1^+/CRE^::Meis2^LoxP/LoxP^ mice challenged by norepinephrine injection failed to adapt compared to WT and Isl1^+/CRE^ mice that presented a large reflex-induced decrease of heart rate (n=6, n=6, n=7 for WT, Isl1^+/CRE^ and Isl1^+/CRE^::Meis2^LoxP/LoxP^ animals respectively; Figure 4B). Similarly, when mice were injected with phenylephrine (Figure 4C), only Isl1^+/CRE^::Meis2^LoxP/LoxP^ mice failed to exhibit the expected heart rate decrease (n=5, n=5, n=6 for WT, Isl1^+/CRE^ and Isl1^+/CRE^::Meis2^LoxP/LoxP^ animals, respectively). These experiments demonstrate that WT and Isl1^+/CRE^ mice responded as expected to these pharmacological compounds, whereas in Isl1^+/CRE^::Meis2^LoxP/LoxP^ mutant mice, the cardio-inhibitory reflex was severely blunted.

**Figure 4:**
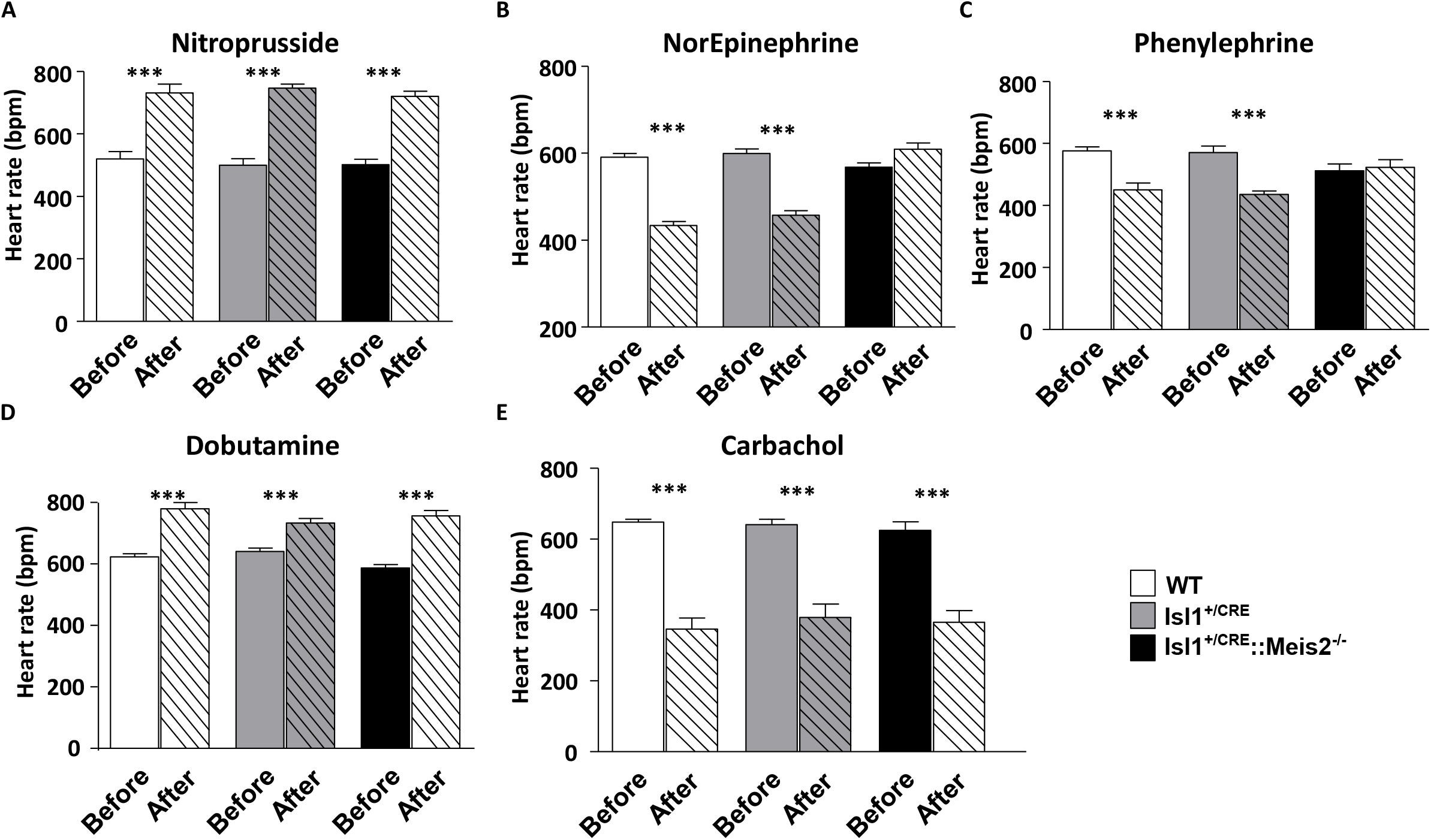
Blunted cardio-inhibitory reflex in Isl1^+/CRE^::Meis2^LoxP/LoxP^ mice. Graphs showing the heart rate adaptation after pharmacologically induced hemodynamic changes in WT, Isl1^+/CRE^ and Isl1^+/CRE^::Meis2^LoxP/LoxP^ vigil mice. Heart rate analyses were done before and after injection of Nitroprusside (A), Norepinephrine (B), Phenylephrine (C), Dobutamine (D) and Carbachol (E). n=6 to 7 mice in each group. \*\*\**p≤0*.*001 vs* before the injection.

Finally, to exclude a default in both cardiac adrenergic and muscarinic signaling pathways in Isl1^+/CRE^::Meis2^LoxP/LoxP^, we injected dobutamine and carbachol known to increase and decrease heart rate respectively by directly acting on cardiac tissues (Figure 4D and E). As expected, in all groups of mice, dobutamine or carbachol injections similarly and significantly increased or decreased respectively the heart rate (*p<0*.*001*). Altogether, these results demonstrate that despite functional adrenergic and muscarinic signaling pathways directly acting on cardiomyocytes, *Meis2* mutant mice were resistant to heart rate adaption when blood pressure was pharmacologically and acutely increased. By contrast, heart rate adapted normally to a pharmacologically induced rapid fall in blood pressure. These results suggest that *Meis2* inactivation interferes with cardio-inhibitory reflexes while cardio-excitatory reflexes remain unaffected.

### Meis2 is required for cardio-inhibitory reflex

Because *Meis2* mutant mice fail to activate cardio-inhibitory reflexes following injection of drugs that induce vasoconstriction, and to exclude a possible failure of norepinephrine of phenylephrine to evoke primary vasoconstriction, blood pressure was simultaneously monitored to heart rate in anesthetized mice before and after supramaximal dose injections of these compounds. We plotted the gain as the variation of heart rate over the variation of blood pressure (ΔHR/ΔBP; Figure 5). Because in the above telemetric experiments no differences were evidenced between control WT and Isl1^+/CRE^ mice, we focused our analysis by comparing WT and Isl1^+/CRE^::Meis2^LoxP/LoxP^ mice only. Under gaseous anesthesia, heart rate was not different between WT and Isl1^+/CRE^::Meis2^LoxP/LoxP^ animals before injection. However, consistent with the low LF/HF ratio in Isl1^+/CRE^::Meis2^LoxP/LoxP^ animals (Figure 2C), their basal systolic, diastolic and mean blood arterial pressures were slightly but significantly lower than in WT mice (*p≤0*.*05*, n=15 in each group; Figure 5A and B).

**Figure 5:**
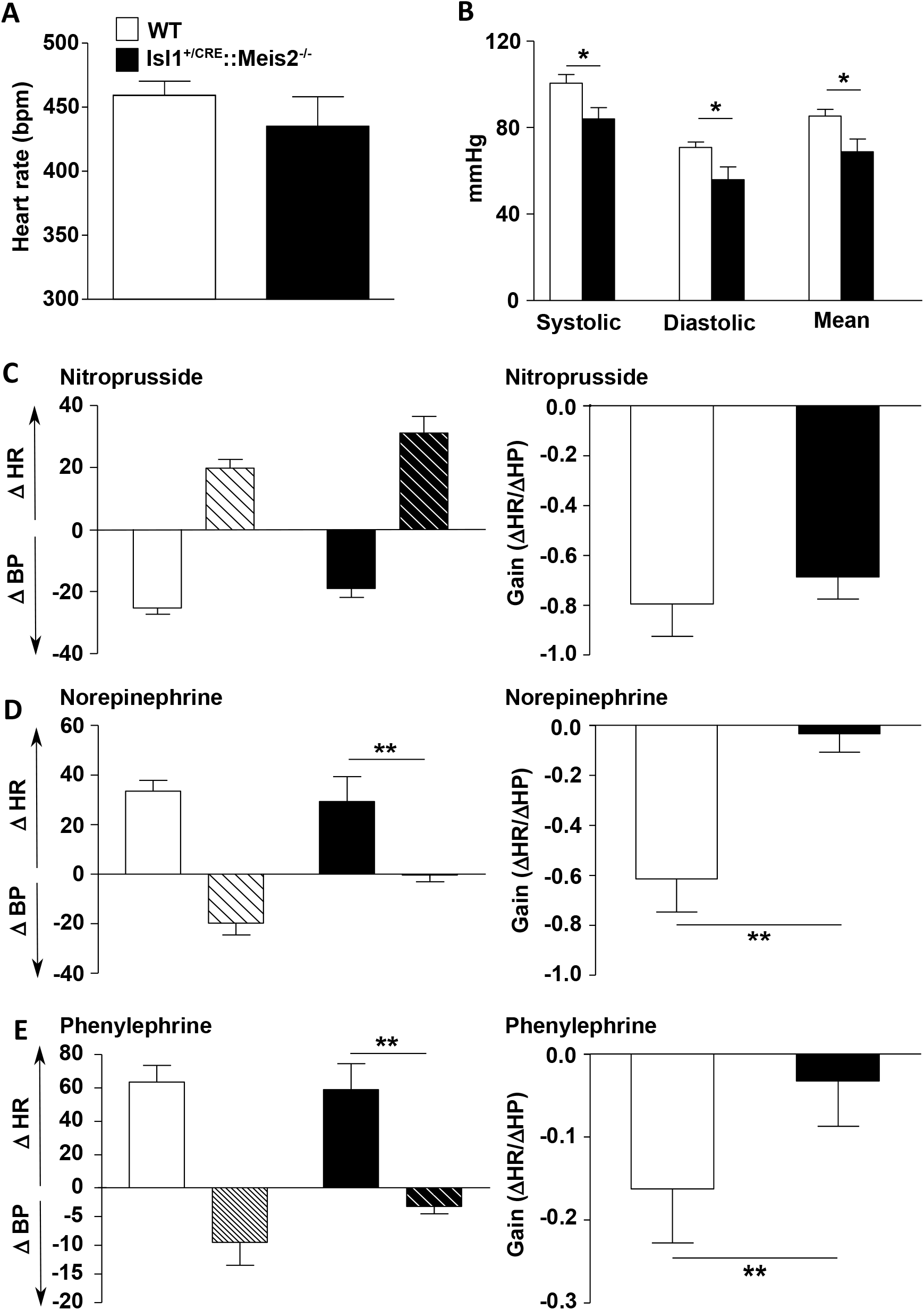
Lack of cardio-inhibitory reflex in Isl1^+/CRE^::Meis2^LoxP/LoxP^ mice. **A**) Graphs showing the measurements of the heart rate (RR) in WT and Isl1^+/CRE^::Meis2^LoxP/LoxP^ anesthetized mice. **B**) Graph showing measurements of blood pressure in WT and Isl1^+/CRE^::Meis2^LoxP/LoxP^ mice. The systolic, diastolic and mean blood arterial pressures at baseline are shown. **C-D**) Graphs showing the variations in mean arterial blood pressure (BP), heart rate (HR) and related gain (ΔHR/ΔBP) following injection of Nitroprusside (**C**), Norepinephrine (**D**) and Phenylephrine (**E**). \**p≤0*.*05*; ** *p≤0*.*01 vs* WT. n=6 to 7 mice in each group.

Following nitroprusside challenge, both WT and Isl1^+/CRE^::Meis2^LoxP/LoxP^ mice exhibited a similar decrease in mean arterial blood pressure leading to a rise in heart rate (Figure 5C, Supplemental figure 1A) that ultimately resumed in a similar ΔHR/ΔBP gain (n=5 per group). When norepinephrine was injected, blood pressure largely increased in both WT (n=6) and Isl1^+/CRE^::Meis2^LoxP/LoxP^ (n=5) mice. However, whereas heart rate decreased in WT mice following norepinephrine injection, it remained stable in Isl1^+/CRE^::Meis2^LoxP/LoxP^ (Figure 6D; Supplemental figure 1B), resulting in an almost null gain (ΔHR/ΔBP) compared to WT. To confirm the absence of cardio-inhibitory response provoked by norepinephrine-induced vasopressor effect, another vasoconstrictor compound was tested. Thus, phenylephrine injection confirmed the lack of baroreflex activation in Isl1^+/CRE^::Meis2^LoxP/LoxP^ mice (n=4 mice per group). While mean arterial blood pressure increased, the heart rate was remained virtually unchanged in Isl1^+/CRE^::Meis2^LoxP/LoxP^ mice (Figure 6E; Supplementary figure 1C).

**Figure 6:**
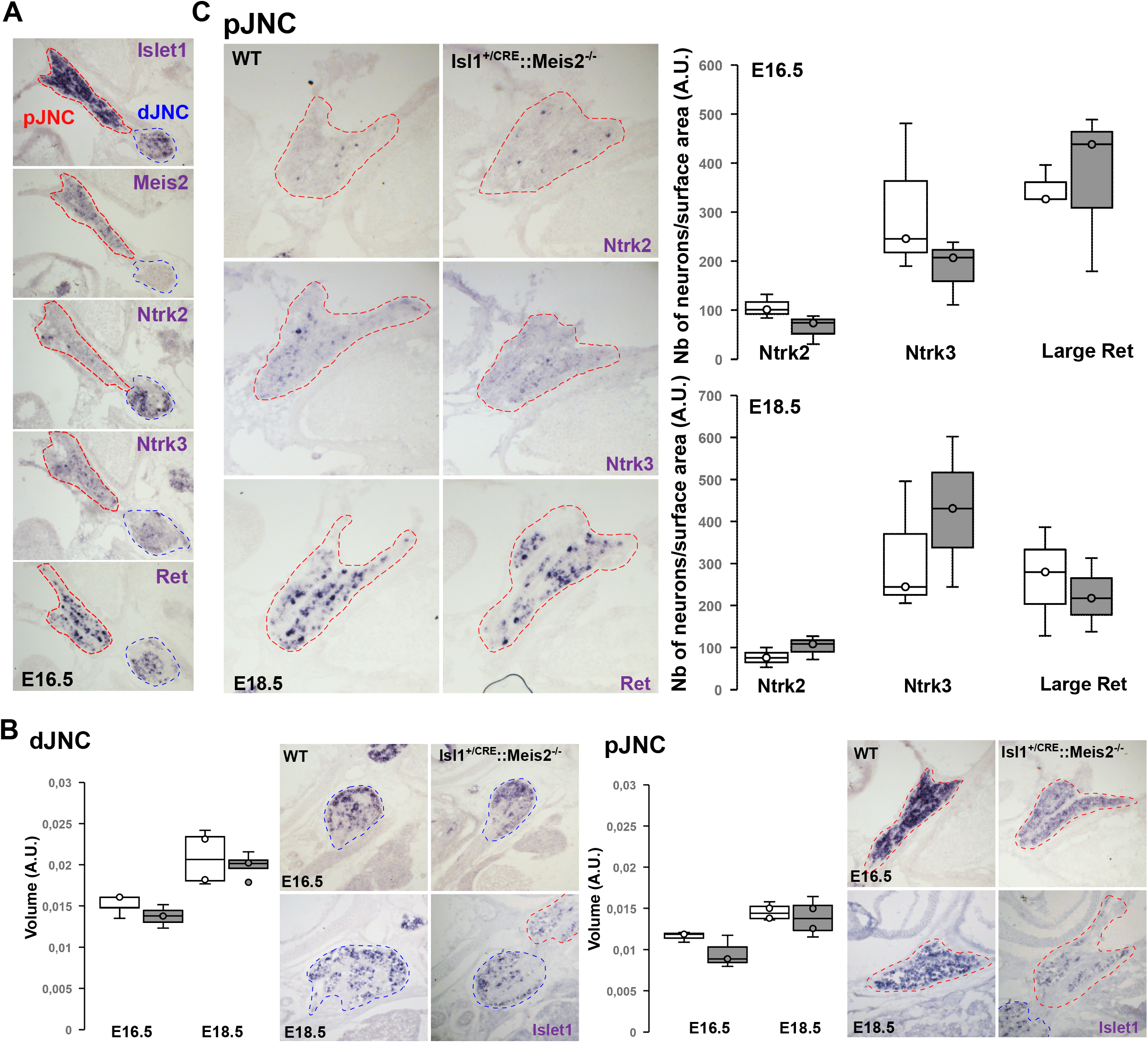
**A**) Expression of Meis2 in the Jugular-Nodose complex of E16.5 mouse embryos. ISH for *Meis2, Islet1*, and other well-known markers for mechanosensitive neurones such as *Ntrk2, Ntrk3* and *Ret* on sagittal sections showed that *Meis2* is expressed by neuronal subpopulations in the proximal Jugular-Nodose complex (pJNC) in a pattern comparable with mechano-sensitive DRG neurons as previously described, but not in the distal Jugular-Nodose complex (dJNC). Red dotted lines delineate the pJNC and blue dotted lines delineate the dJNC. **B**) Size Measurements of the pJNC and dJNC following ISH for Islet1 showed no change in the size of the ganglia at E16.5 and E18.5 of Isl1^CRE^::Meis2^LoxP/LoxP^ embryos compared to WT embryos (n=3 for each genotype). Images are representative of the different ISH staining in the pJNC and dJNC. **C**) Graphs showing the counting of Ntrk2, Ntrk3 and large Ret-positive neurons in the pJNC at E16.5 and E18.5. No changes were evidenced in the number of these neurons in Isl1^+/CRE^::Meis2^LoxP/LoxP^ embryos compared to WT (n=3 for each genotype). Images are representative of the different ISH staining in the pJNC.

These results demonstrate that *Meis2* inactivation in *Isl1*-expressing cells impedes the cardio-inhibitory reflex that in normal conditions preserves cardiovascular homeostasis, whereas its counterpart, the tachycardic reflex, is normal. Both WT and Isl1^+/CRE^::Meis2^LoxP/LoxP^ mice presented functional adrenergic and muscarinic signaling pathways and developed the expected responses when blood pressure is challenged by vasodilator and vasoconstrictor drugs^41^, strongly supporting that heart and artery contractile activities are normal. Collectively our results suggest that rather than alteration of heart and/or arteries intrinsic functionality, the normal functioning of peripheral sensory neurons involved in this reflex is impaired.

Our mouse model strikingly resembles a recently reported mouse model conditionally targeting Piezo channels in mechanosensitive neurons of the nodose ganglia^42^. In this mouse model, genetic ablation of both Piezo1 and Piezo2 mechanosensitive channels in the nodose and petrosal sensory ganglia abolished vasopressor (phenylephrine)-induced bradycardic effect and aortic depressor nerve activity. This results in the abolishment of baroreflex (ΔHR/ΔBP) and an abnormal adaptation of heart rate to blood pressure modification. Moreover, pulse BP measurement revealed a great variability of beat-to-beat control of heart rate at baseline. We and others’ single cells RNAseq (scRNAseq) analyses have previously reported Meis2 expression in peripheral sensory neurons^36,43–46^. In spinal DRG sensory neurons, Meis2 is expressed by subclasses of mechanosensitive neurons mediating touch perception^36,43–45^, and we showed, using the Isl1^+/CRE^::Meis2^LoxP/LoxP^ mouse strain, that specific post-mitotic *Meis2* inactivation in those neurons impaired their electro-physiological responses to mechanical stimuli and strongly altered evoked-touch perception. In a scRNAseq performed on vagal sensory neurons^46^, *Meis2* expression is reported in distinct neuronal subpopulations that also express Piezo2 (Supplementary Figure 2; https://ernforsgroup.shinyapps.io/vagalsensoryneurons/), and that are molecularly similar to mechanosensitive neurons of the DRG. We thus investigated, in E16.5 mouse embryos, *Meis2* expression by *in situ* hybridization together with several classical identity markers for these mechanosensitive subclasses (Figure 6A). A strong signal for *Islet1* mRNA was observed in the jugular-nodose complex (JNC) including the proximal (pJNC) and the distal (dJNC) parts of the complex. *Meis2* mRNA expression was mostly restricted to neuronal subpopulations located in the proximal complex together with subpopulations of neurons expressing Ntrk2, Ntrk3 or Ret. Measures of dJNC and pJNC volumes on consecutive sagittal sections of E16.5 and E18.5 WT and Isl1^+/CRE^::Meis2^LoxP/LoxP^ embryos however did not show any significant differences (Figure 6B). Finally, the numbers of Ntrk2-, Ntrk3 and large Ret-expressing neurons were not significantly different between the pJNC of E16.5 and E18.5 WT and Isl1^+/CRE^::Meis2^LoxP/LoxP^ embryos (Figure 6C).

## Discussion

In this work, we showed that specific inactivation of Meis2 TF in Isl1-expressing cells in mice severely impaired inhibitory baroreflex function independently of any developmental cardiac malformations or contractility defects of the heart and cardio-vascular system. In addition, the *Meis2* expression in subclasses of vagal neurons that we and others reported and that are predicted to have proprioceptive and mechanosensitive properties^46^, together with the recent demonstration that Piezo2-positive vagal neurons are essential for the cardiac baroreflex^42^, strongly suggest that *Meis2* inactivation in those neurons could be responsible for the blunted inhibitory cardiac reflex we report. In this scenario, *Meis2*-expressing mechanosensitive neurons, including those from the JNG and possibly the DRG whose function is to sense stretch induced by artery and/or heart deformations fail to properly encode the information necessary to trigger the normal inhibitory baroreflex feedback. Thus, our model reinforces current hypotheses on alterations of primary sensory neurons function in ASD disorder^47^, and underline the importance of conditionally targeted mouse models to disentangle intermingled and complex phenotypes found in human mutants.

The baroreflex is a classical and complex mechanism that coordinates adaptive cardio-vascular tone through both autonomic and sensory components^48^. Elevated blood pressure promptly triggers a compensatory decrease in cardiovascular output to maintain body and brain blood pressure within homeostatic ranges^49^. There is no real consensus about the sensory neuron subtypes involved. They are commonly called baroreceptors, display mechanosensitive properties and project to precise locations on arteries where they sense arterial wall distortion. This arterial baroreceptor reflex system plays a dominant role in preventing short-term wide fluctuations of arterial blood pressure, as recurrently demonstrated in experiment where arterial baroreceptor denervation leads to an increase of the beat to beat variability of blood pressure and related heart rate^50^.

The baroreflex is associated with some pathological conditions^48,51^, but only recently, imbalance of cardiac autonomic regulation in patients with intellectual disabilities and ASD is emerging^7–19^. However, the origins of dysautonomia in ASD is still unclear and somehow controversial with highly variable profiles depending on the studies. Many studies report that ASD patients present a higher heart rate, and that exposure to external stimuli leads to a blunted heart rate response compared to healthy subjects^8^. HR is increased in ASD patients compared to control due to a lower parasympathetic activity^17,18^, but other reports revealed on the contrary an increased parasympathetic activity^15^. Moreover, intermittent neuro-cardiovascular autonomic dysfunction affecting heart rate and blood pressure was also over-represented in ASD^52,53^.

Interpretation of results in human patients has proven complicated due to the genetic variability causing the different syndromes and the combinatory effect of multiple affected organs other than the nervous system. In most investigations related to cardiac autonomic regulation and HRV analysis in ASD, patients are rigorously matched in age and gender but cohorts usually do not take into account the genetic basis of the diagnosed ASD. Indeed, the large number of neurodevelopmental genes supporting ASD symptoms, but also the variability of the symptoms accompanying different mutations within the same gene could account for discrepancies between studies. A good example linking gene dosage effect to the severity of phenotypic manifestation is Rett syndrome. Rett syndrome is associated with *MECP2* gene mutations, but the type of mutation, *i*.*e*. loss-of-function, gene duplication or triplication, and the degree of mosaicism for these mutations within cell types lead to highly heterogeneous phenotypic manifestations and clinical presentation ranging from microcephaly to normal brain size, shortened lifespan or not^54^. Nevertheless, studies have shown modified autonomic function both in children and adult ASD patients overall characterized by a lower autonomic nervous activity than healthy subjects.

The genetic links between ASD and congenital heart malformation in humans also prevent unmasking deleterious effects on cardiac autonomic regulation in ASD full knockout mouse models^55^. Recent advances in the understanding of the biology of the MEIS family of TFs and their well-known partners PBX members emphasized their essential contribution to cardiac morphogenesis and physiology. In humans, non-synonymous variants for *PBX1, PBX2, PBX3, MEIS1* and *MEIS3* have been identified in patients with congenital cardiac defects^56^, and humans carrying *MEIS2* mutations present cardiac septal defects^27,28^. Similar phenotypes are also described in full-knockout models for those genes^33,34^. In mouse, genetic ablation of *Pbx1-3* at specific developmental stages lead to heart malformations^34^. *Pbx1* deficiencies results in persistent truncus arteriosus, whereas *Pbx2* and *3* inactivation leads to Pbx1 haploinsufficiency with overriding aorta, ventricular septal defect, and bicuspid aortic valves^34^. *Meis1* and *Meis2* mutant mice also exhibit cardio-vascular and septal defects^31,33,34,57,58^.

Surprisingly, our Doppler-echocardiography investigations did not reveal any heart morphological or contractile defects. This might be due to a later *Meis2* inactivation in cardiac neural crest compared to the AP2α-IRES-Cre strain used by others^33^, at a time when *Meis2* is no longer required. Given that both genes are involved in heart morphogenesis, this might also result from redundant *Meis1* and *Meis2* activities within the timeframe of our genetic ablation. Nonetheless, we previously showed that *Meis1* ablation-induced septal defect depends on the CRE strain used for neural crest gene ablation. *Meis1* inactivation in early neural crest resulted in septal defects, but *Meis1* inactivation in late neural crest did not produce contractile and morphological defect^31^.

We also showed that at baseline conditions, *Meis2* mutant mice do not present symptomatic heart rhythm disturbance such as major sinus pause or arrest or atrial/ventricular ectopic beats. Instead, a large variability in sinus rhythm confirmed by high HRV, without brady- or tachycardia, was observed. A profound sinus node dysfunction in *Meis2* mutant is thus unlikely. We further demonstrated that the large beat-to-beat variability in *Meis2* mutant mice results from a dysregulation of the sensory-autonomic control of cardiac rhythm. We identified a lower sympatho-vagal activity at baseline reflected by the decrease LF/HF ratio that could also explain the low mean arterial pressure observed in mutant mice. When using drugs that rapidly and robustly modify blood arterial pressure, we unmasked a sensory-autonomic dysregulation characterized by a blunted cardio-inhibitory reflex. Surprisingly, only the cardio-inhibitory baroreflex was affected, but the sympathetic activation following a fall in blood pressure was maintained although both vasoconstriction and vasodilation could be pharmacologically elicited.

According to the vagal dominance in the beat-to-beat baroreflex adjustment of HR and blood pressure, and the increased variability during baroreceptor denervation reported in several animal models^50^, we suggest a defective baroreceptor related-vagal pathway induced by *Meis2* inactivation. Moreover, because *Meis2* is not expressed in sympathetic neurons^31^, along with the observation that basal mean heart rate is unaffected and the cardio stimulatory reflex seems to be unaltered, we can exclude that sympathetic nerves were affected by *Meis2* deletion.

Instead, we conclude that *Meis2* inactivation interferes with the sensory component of the vagal-mediated baroreflex. First, because of the possible *Meis2* recombination sites following CRE activity when using the Isl1^CRE^ strain. The LIM-homeodomain TF Isl1 is expressed by several neural and non-neural tissues both during embryonic development and postnatal life amongst which the peripheral and central nervous systems, the pancreas, the heart and the pituitary gland^59–63^. However, interaction of tissues other than the nervous system with the baroreflex is unlikely. Beside sensory and autonomic peripheral neurons, specific CNS neuronal populations express Isl1 including spinal motor neurons, retinal ganglion cells, hypothalamic, central amygdala and striatal neurons^30,64–70^. We therefore cannot fully exclude that *Meis2* recombination also occurs in some of these central neuronal populations that somehow participate in the autonomic imbalance we report in Isl1^+/CRE^::Meis2^LoxP/LoxP^ mice.

Secondly, the large literature linking sensory neurons to baroreflex and our recent finding that *Meis2* is necessary to normal functioning of peripheral mechanosensitive neurons place Meis2-expressing peripheral sensory neurons in best position to support the lack of cardio-inhibitory reflexes in these mice. Beside the autonomic system that includes parasympathetic and sympathetic efferents and control heart and blood vessels contractility, peripheral sensory innervation of the heart is of dual origin^71,72^. Anatomically, sensory fibers originate from vagal neurons located in the jugular-nodose complex and run through the vagus and the inferior cardia nerve^72^. Afferent sensory fibers sense local target organs activities such as tissue tension and send the information to higher brain structures in order to elaborate an adapted response. Although most afferent and efferent information to the heart navigates through the vagus nerve^46,49^, there is evidence that DRG sensory neurons are also involved, in particular for cardiovascular reflexes^72–74^. Retrotracing experiments in cats, dogs and rats injected at different locations in the heart, coronary artery or in the inferior cardiac nerve labelled neurons in the DRG indicating that heart and arteries also receive afferent sensory fibers from the DRG^75–80^. In addition, molecular characterization of these neurons revealed that they express a range of markers compatible with the identity of several subclasses of DRG sensory neurons^79,81,82^.

Vagal neurons have been studied for very long time, but the knowledge and understanding of the precise identities and physiological functions of the different subpopulations of vagal sensory neurons remain fragmented mainly because of the lack of molecular knowledge and tools to specifically target them. Nerve sectioning experiments combined with the mixed nature of the vagus nerve also impedes full interpretation on their precise function. Cranial ganglia contributing to the vagal nerves are multiple and arise from the neural crest derived jugular ganglia and the placode-derived nodose and petrose complex that eventually merge during embryogenesis^83^. The molecular characteristics of these primary sensory neurons have only been very recently elucidated and showed that nodose and jugular neurons are molecularly fundamentally different with jugular neurons sharing many features with somatosensory DRG neurons^46^. In this scRNAseq study^46^, Meis2 was detected in 2 of the 18 nodose neuron clusters, and in 4 of the 6 jugular neurons clusters with relatively high expression in clusters displaying a molecular profile similar to myelinated DRG neurons involved in gentle touch. Functional classification of nodose clusters predicted *Meis2* expressing populations to have DRG proprioceptive-like features. Interestingly, in this database (https://ernforsgroup.shinyapps.io/vagalsensoryneurons/), the mechano-sensitive Piezo2 channel recently shown to be involved in baroreflex^42^ was coexpressed in all Meis2-expressing clusters (Supplementary Figure 2). Thus, mechanosensitive neurons of the DRG and of the Jugular-Nodose complex are molecularly highly similar.

In our mouse model, we could not evidence any neuronal loss of vagal neurons of the JNG, suggesting that *Meis2* inactivation does not affect neuronal survival or identity as demonstrated by the normal expression of Ntrk2, Ntrk3 and Ret. These results are in line with our previous studies on *Meis1* or *Meis2* inactivation in different types of peripheral neurons^31,36^. When *Meis1* is specifically inactivated in sympathetic neurons, distal innervation of target organs, including the heart, is compromised, but early sympathetic specification is unaffected^31^. More strikingly, using the very same mouse strain as in the present study, we found that mechanosensitive neurons of the DRG that normally express *Meis2* failed to fully differentiate and to elaborate complex distal peripheral sensory terminals mediating touch sensation in the skin^36^. In both mouse models, these peripheral innervation defects result in physiological consequences: mice lacking *Meis1* in sympathetic neurons display severe chronotropic incompetence due to sympathetic dysfunction and mice lacking *Meis2* in DRG mechanosensitive neurons have impaired touch sensations. Using an another conditional *Meis2* strain, Machon et al. inactivated *Meis2* in the neural crest, including the neural crest-derived cranial sensory ganglia encompassing trigeminal (V), facial (VII) and vestibulocochlear (acoustic) nerves (VIII)^33^. Although the authors did not thoroughly detail their findings, most neural crest-derived cranial ganglia were reported to be present, but nerves exiting the ganglia seemed less numerous and less ramified than in WT embryos as seen by whole mount neurofilament staining. However, in this study, the physiological consequences have not been investigated.

To conclude, although we could not unambiguously demonstrate that Meis2 expressing vagal and/or DRG neurons are directly responsible for the blunted autonomic response and the lack of baroreflex in Isl1^+/CRE^::Meis2^LoxP/LoxP^ mice, our study clearly showed that our genetically modified animal model is a very appropriate tool to study autonomic dysregulation independently of cardiac remodeling.

## Material and Method

### Animals

All protocols complied with Directive 2010/63/EU of the European Parliament and the Council of 22 September 2010 for the protection of animals used for scientific purposes and Ethics committee for animal experiments, Languedoc Roussillon, C2EA -36 (agreement: B34-172-38; project APAFIS#11026). All efforts were made to minimize animal suffering during the experiment and to reduce the number of animals used by performing echocardiography and ECG recording in the same animal when possible, and for pharmacological dosing (2 days of washing period). Animals were housed in a temperature-regulated room (12 h day/12 h night cycle) with ad libitum access to food and water. Protocols were only conducted by trained and authorized experimenters.

The two genetically modified strains of mice used in this study have previously been reported^36,37^. Mice of three different genotypes were analyzed in the present study for *in vivo* experiments, wild-type (WT, n=15, weight = 22.9±1.8 g), Isl1^+/CRE^ (n=10, weight = 20.2±1.9 g) and Isl1^+/CRE^::Meis2^LoxP/LoxP^ strain (n=15, 21.4± 1.7), aged 3 months, randomly assigned (equal proportion of males and females).

### Echocardiography

Mice were anesthetized with 1.5% isoflurane in 100% oxygen to reach comparable heart rate and placed on a heating table in a supine position. Body temperature was monitored through a rectal thermometer to be maintained at 36–38° C and ECG was recorded all along the echocardiographic procedure with limb electrodes. Ejection fraction (EF%) and fractional shortening (FS%) were calculated from the left ventricular internal diameters (LVID) on M-mode measurements at the level of papillary muscles in a parasternal short-axis two dimensional view using Vevo 2100 (VisualSonics, FujiFilm, Netherlands). To better consider left ventricular morphology and possible outflow tract remodeling or malformation, EF was also calculated from a B mode parasternal long axis view (EF% B-mode) by tracing endocardial end-diastolic and end-systolic borders to estimate left ventricular volumes, and the endocardial fractional area change (FS%) on a parasternal short-axis view at papillary muscle level was similarly measured. Mitral flow was recorded by a pulsed-wave Doppler sampling at the tips of the mitral valves level from the apical four-chamber view. Peak early (E) and late atrial contraction (A) mitral inflow waves velocities were measured and the ratio E/A was calculated. Pulsed-wave Doppler of the ascending aortic blood flow was recorded permitting measurements of the velocity time integral (AoVTI). All measurements were quantified and averaged for three cardiac cycles as previously done^31^.

### ECG in conscious mice

Recording: Electrocardiogram (ECG) were monitoring by telemetry system on vigil unrestrained mice. After a preanesthetic (physical) evaluation, the transmitter (PhysioTel, ETA-F10 transmitter) was inserted in mice subcutaneously along the back under general anesthesia (2% inhaled isoflurane/O2, Aerrane, Baxter, France) coupled with local anesthetic (lidocaine 0.5 %), and two ECG electrodes were placed hypodermically in the region of the right shoulder (negative pole) and toward the lower left chest (positive pole) to approximate lead II of the Einthoven surface ECG. During procedure, respiratory and cardiac rate and rhythm, adequacy of anesthetic depth, muscle relaxation, body temperature, and analgesia were monitored to avoid anesthesia-related complications. Post-operating pain was considered during one-week post-implantation period and buprenorphine (0.3 mg.kg-1 sc) could be done at case per case. ECG monitoring were performed 2 weeks after recovery from surgery in the home cage with a signal transmitter-receiver (RPC-1) connected to a data acquisition system (Ponemah system, Data Sciences International, Saint Paul, USA). Data were collected continuously over 24 h at a sampling rate of 2000 Hz as previously^84^.

ECG recording were also performed after pharmacological injection of Nitroprusside (2.0 mg/k-1g ip, NaCl, 0.9%), Norepinephrine (2.5 mg/kg-1 ip, NaCl 0.9%), Phenylephrine (2.5 mg.kg-1 ip, NaCl 0.9%), Dobutamine (1 mg/kg-1 ip) and Carbachol (0.5 mg.kg-1 ip, NaCl 0.9%) according to literature. All molecules were purchased at Sigma-Aldrich (France) and diluted in NaCl sterile solution (Aguettant, France). Sodium Nitroprusside is a major vasodilator by acting on NO release and induces a pronounced reflex tachycardia. Norepinephrine is the neurotransmitter released by postganglionic neurons of sympathetic system (α1 and β1 adrenergic receptor agonist) inducing a major hypertension followed by reflex-bradycardia. Phenylephrine is a specific α1-adrenergic receptor agonist, increasing peripheral resistance and blood pressure that precipitates in sinus bradycardia due to vagal reflex. Dobutamine is a sympathomimetic, mainly through β1 adrenoreceptors activation leading to a rapid raise of heart rate by acting directly on cardiomyocytes. Carbachol is a nonselective muscarinic receptor agonist leading to profound direct bradycardia.

ECG waveforms analysis: Continuous digital recordings were analyzed off line after to be digitally filtered between 0.1 and 1,000 Hz. ECGs during nocturnal periods (12-hours) and during pharmacological testing were analyzed with Ponemah software using template automatic detection, secondly validated by an operator. The mean RR interval and the mean PR, QRS, QT durations were calculated. The QT interval was defined as the time between the first deviation from an isoelectric PR interval until the return of the ventricular repolarization to the isoelectric TP baseline from lead II ECGs^38^. Presence of potential ectopic beats were scanned by hand. Specific to pharmacological testing, parameters were measured 2-hours before injection and compared to value obtained between 2 to 10 min post-dosing at maximum responses as previously reported^38,85^. A minimum of 15 complexes were used for analysis and averaged.

HRV analysis: the autonomic nervous system adapts continuously heart rate to metabolic needs, inducing beat to beat heart rate variability by modifying the automatic sinus activity through a complex interplay of the ortho-sympathetic and parasympathetic (or vagal) systems. Time-and frequency domain indices of HRV are the standard parameters to evaluate ANS activity as well in clinics as in fundamental research. Total variability was assessed with the standard deviation of all normal RR intervals (SDNN) in the time domain. HRV was also evaluated by power spectra analysis (ms^-2^) using the fast Fourier transformation (segment length of 2048 beats, linear interpolation with resampling to a 20-Hz interbeat-time series and Hamming windowing). The cut-off frequency ranges for the low frequency (LF: 0.15–1.5 Hz) and high frequency powers (HF: 1.5–5 Hz) were chosen according to those used in the literature. As in humans, the low frequency reflects a complex interaction between sympathetic and parasympathetic ways that modulates heart rate including baroreflex function^86^. The efferent vagal activity rests the major contributor to the HF component, as seen in clinical and experimental observations of autonomic maneuvers such as electrical vagal stimulation, muscarinic receptor blockade, and vagotomy. Thus, as previously performed^31,85^, the cardiac sympathetic and spontaneous baroreflex activities were assessed from LF, the vagal activity was assessed from HF and the LF/HF ratio, conjointly with the mean values of HF and LF power, was used to assess sympatho-vagal activity on heart rhythm.

### BP and HR recordings under anesthetized conditions

To determine the origin of response failure of some molecules observed by telemetry, we recorded blood pressure coupled to heart rate change in anesthetized animal (2% inhaled isoflurane/O2, Aerrane, Baxter, France) using Powerlab system and LabChart software (Blood pressure module; ADInstrument Ltd, France). A Millar Mikro-Tip® pressure catheter is introduced in carotid to assess diastolic, systolic and mean arterial blood pressures. In parallel, ECG was recorded using lead II Einthoven derivation. Parameters were measured in baseline conditions and after injection of Nitroprusside (2.0 mg/k-1g ip, NaCl, 0.9%), Norepinephrine (2.5 mg/kg-1 ip, NaCl 0.9%) and Phenylephrine (2.5 mg.kg-1 ip, NaCl 0.9%). Parameters were measured and averaged during maximal response, on fifteen complexes. The delta of heart rate and delta of mean BP were calculated. The Gain (Delta HR / Delta BP) was done and reflects cardiovascular adaptation during pharmacological dosing through baroreflex.

### In situ Hybridization

RNA probes used in the study and in situ hybridization procedures have been reported previously^31,36^. Briefly, tissues were collected and fixed in 4% paraformaldehyde/PBS overnight at 4°C and incubated overnight at 4°C for cryopreservation in increasing sucrose/PBS solutions (10 to 30% sucrose). After snap freezing in TissueTek, embryos were sectioned at 14-µm thickness and stored at -20°C until use. Before hybridization, slides were air dried for 2-3 hours at room temperature. Plasmids containing probes were used to synthesize digoxigenin-labeled or fluorescein-labeled antisense riboprobes according to supplier’s instructions (Roche) and purified by LiCl precipitation. Sections were hybridized overnight at 70°C with a solution containing 0.19 M NaCl, 10 mM Tris (pH 7.2), 5 mM NaH2PO4*2H2O/Na2HPO4 (pH 6.8), 50 mM EDTA, 50% formamide, 10% dextran sulphate, 1 mg/ml yeast tRNA, 1XDenhardt solution and 100 to 200 ng/ml of probe. Sections were then washed four times for 20 minutes at 65°C in 0.4X SSC pH 7.5, 50% formamide, 0.1% Tween 20 and three times for 20 minutes at room temperature in 0.1 M maleic acid, 0.15 M NaCl and 0.1% Tween 20 (pH 7.5). Sections were blocked 1 hour at room temperature in presence of 20% goat serum and 2% blocking agent (Roche) prior to incubation overnight with AP-conjugated anti-DIG-Fab-fragments (Roche, 1:2000). After extensive washing, hybridized riboprobes were revealed by performing a NBT/BCIP reaction in 0.1 M Tris HCl pH 9.5, 100 mM NaCl, 50 mM MgCl2 and 0.1% Tween 20. Wide field microscopy (Leica DMRB, Germany) was used to take the images.

### Statistical analysis

All values are expressed as means□±□SEM. For data from more than two experimental groups, one-way or two-way ANOVA was used to assess group means followed by the Bonferoni post-Hoc test. Paired comparisons were made if needed. P ≤ 0.05 was taken to denote statistical significance.

## Figures legends

**Supplementary Figure 1:** Representative traces showing the mean blood arterial pressure (MAP) and related heart rate recording during pharmacological injection of Nitroprusside (A), Norepinephrine (B), and Phenylephrine (C) in anesthetized WT and Isl1^+/CRE^::Meis2^LoxP/LoxP^ mice.

**Supplementary Figure 2:** Graphs showing Islet1, Meis2 and Piezo2 expression extracted from scRNAseq of jugular and nodose ganglia (Data were extracted from https://ernforsgroup.shinyapps.io/vagalsensoryneurons/). Note that Islet1 is expressed by all neuronal populations ensuring recombination in all vagal neurons. Meis2 is expressed by 5 out of the 6 clusters of jugular neurons and in 2 of the 18 clusters of nodose neurons. In all those clusters, neurons also express Piezo2.

## Acknowledgment

The authors thank Patrick Carroll for critical reading of the manuscript.

## Notes

### Competing Interest Statement

The authors have declared no competing interest.

