## Supplementary figures and images for "The autism-associated *Meis2* gene is necessary for cardiac baroreflex regulation in mice"

### Supplementary 1

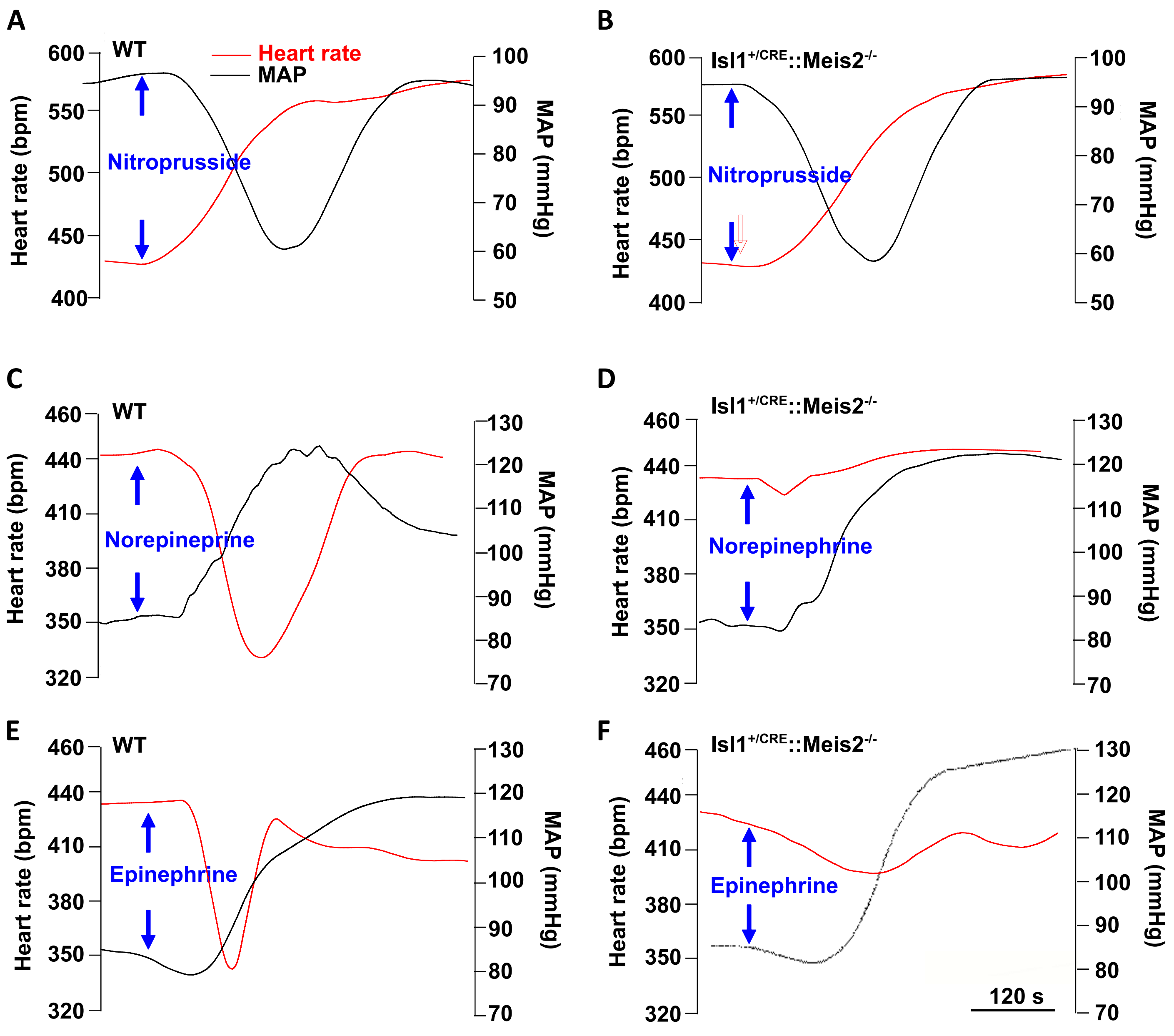


**Figure Supplementary 1**

### Supplementary 2

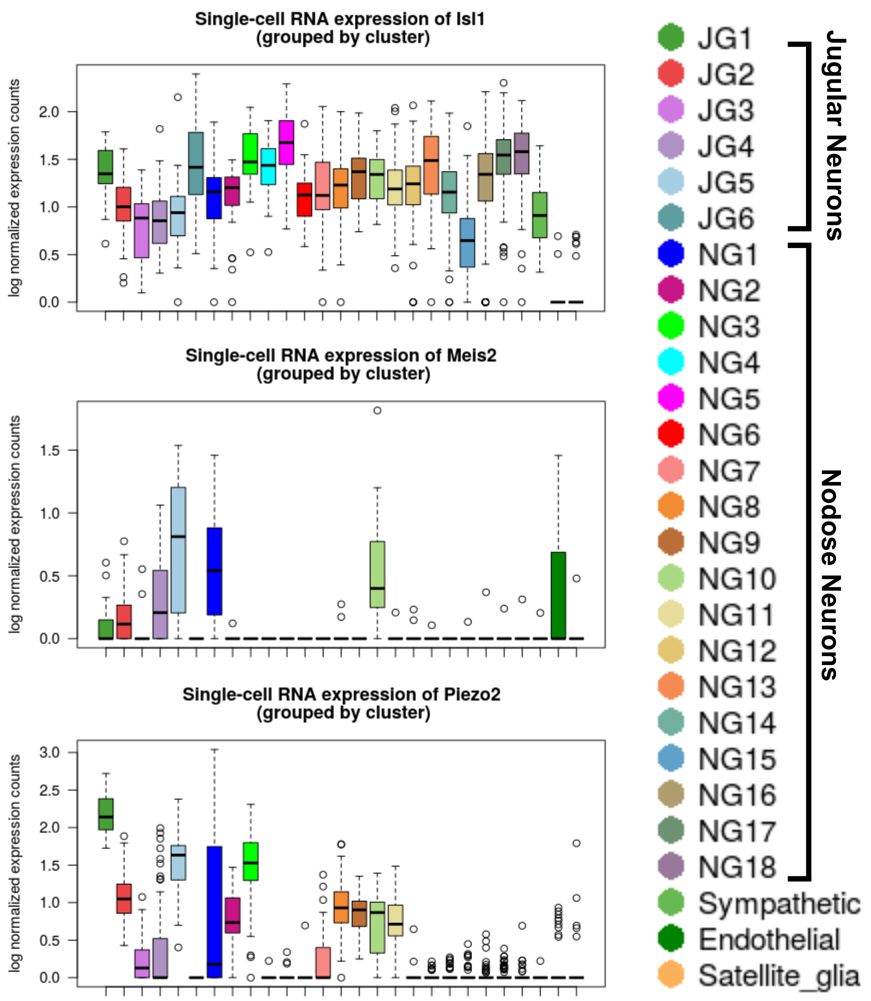


**Figure Supplementary 2**
